# Influenza A Virus Ribonucleoproteins Form Liquid Organelles at Endoplasmic Reticulum Exit Sites

**DOI:** 10.1101/410373

**Authors:** Marta Alenquer, Sílvia Vale-Costa, Ana Laura Sousa, Temitope Akhigbe Etibor, Filipe Ferreira, Maria João Amorim

**Author notes:** Equal contribution (Ana Laura Sousa, Temitope Akhigbe Etibor).

## Abstract

Influenza A virus has an eight-partite RNA genome that during viral assembly forms a supramolecular complex containing one copy of each RNA. Genome assembly is a selective process driven by RNA-RNA interactions and is thought to lead to discrete punctate structures scattered through the cytosol. Here, we show that contrary to the accepted view, formation of these structures is not dependent on RNA-RNA interactions among distinct viral ribonucleoproteins (vRNPs), as they assemble in cells expressing only one vRNP type. We demonstrate that these viral inclusions display characteristics of liquid organelles, segregating from the cytosol without a delimitating membrane, dynamically exchanging material, deforming easily and adapting fast to hypotonic shock. We provide evidence that they develop close to the Endoplasmic Reticulum Exit Sites (ERES), being dependent on continuous ER-Golgi vesicular cycling. We show that viral inclusions do not promote escape to interferon response, and propose that they facilitate selected RNA-RNA interactions in a liquid environment of concentrated vRNPs.

## MAIN TEXT

Influenza A infections are serious threats to human health causing annual epidemics and occasional pandemics^1^. The virus contains an eight-partite RNA genome, and each segment is encapsidated as an individual viral ribonucleoprotein (vRNP) complex. vRNPs are composed of single-stranded negative-sense RNA, with base paired terminal sequences originating a double stranded RNA portion where the trimeric RNA-dependent RNA polymerase (RdRp), composed of PB1, PB2 and PA, binds. The remaining sequence attaches several copies of unevenly-bound nucleoprotein (NP)^2^. The advantages of having a segmented genome are evident for viral evolution^3^ and for better gene expression control^4^, but increase the complexity of the assembly of fully infectious virions^5-8^.

Viral assembly occurs at the plasma membrane and, in 80% of the cases, 8 distinct vRNPs are packaged selectively into a budding membrane^9^. Seminal work established the requirement of cis-acting and intersegment RNA-RNA interactions for the formation of this supra-molecular complex (reviewed in^5-8^). However, it is under debate if vRNPs reach the plasma membrane already as complete genome bundles.

Upon exiting the nucleus, where they replicate, vRNPs accumulate around the microtubule organizing centre^10^ and, subsequently, distribute throughout the cytoplasm concentrating in discrete puncta that enlarge as infection progresses^10-14^. Each puncta accommodates different vRNP segments with the diversity in vRNPs increasing proportionally to the proximity of the plasma membrane^11,13^. Such observation led to the proposal that genome assembly preceded vRNP packaging in budding virions by a process intimately linked with the formation of the referred vRNP hotspots^6,11,13-15^. Studies on the biogenesis of vRNP hotspots showed that their formation requires the cellular GTPase Rab11^10,12,16,17^. In uninfected cells, Rab11 is the master regulator of the endocytic recycling compartment (ERC), one of the systems the cell uses for delivering endocytosed material, as well as specific cargo from the *trans-Golgi-Network* (TGN), to the cell surface^18^. The process is very well-described for uninfected cells, with Rab11-GTP regulating ERC transport by recruiting molecular motors, tethers and SNARES to respectively drive, dock and fuse vesicles to the plasma membrane^18^. Despite initial reports that the functional role of Rab11 was to deliver vRNPs to the cell surface^10,12,17,19^, accumulating evidence analyzing Rab11 sub-cellular localization and non-abundancy in virions^14,16,20^, binding partners^14^ and host transferrin recycling^14,21^ strongly indicates that Rab11 is redirected and its function is impaired during IAV infection. In fact, it was demonstrated that vRNPs outcompeted Rab11 effectors for Rab11 binding, rendering the recycling process sub-optimal^14^. Further corroborating the scenario that Rab11 pathway is impaired with infection, a recent publication showed that Rab11 was re-routed to the ER during IAV infection^16^. In addition, using correlative light and electron microscopy, vRNP hotspots were shown to concentrate clustered vesicles positive for Rab11, surrounded by electron dense material^14^.

The formation of vRNP hotspots was postulated to be dependent on the establishment of sequential RNA-RNA interactions occurring as Rab11 vesicles transporting vRNPs collided^6,11,13,22,23^. However, the impaired ERC hypothesis above mentioned, argues against the requirement for RNA-RNA interactions in the formation of vRNP hotspots and challenges the IAV assembly model proposed. Nevertheless, the existence of vRNP/Rab11 hotspots indicates segregation from the cytosol in foci that are not delimitated by membranes, although they contain numerous remodeled membranes inside^14,15^. Sub-organelles not delimitated by membranes are abundant in the viral world and are known as viroplasms, viral factories, aggresomes or virosomes, to indicate sites of viral replication^24-26^. Viruses can also form viral inclusions and these are sites of accumulation of viral proteins, nucleic acid and selected host proteins and can include viral factories or not^24,25^. Given this definition, IAV vRNP hotspots could be re-classified as viral inclusions. The most notable cases of electron dense aggregated material in the cytosol (not delimited by membranes) are found in cells infected by viruses of DNA (*Poxviridae, Iridoviridae, Asfaviridae*), of dsRNA (*Reoviridae*) and of negative-sense RNA genome (*Paramyxoviridae, Rhabdoviridae, Filoviridae*)^24-29^. Formation of factories is associated with remodeling of host membranes and/or cytoskeleton to orchestrate sophisticated platforms for viral replication and/or for escaping host immune recognition^26^. However, several questions remain unclear relative to the internal organization and biophysical properties of these cellular condensates. Resolving these questions for IAV will help to understand the rules and physical properties of organizing cellular matter into membraneless organelles in the virus world and identify their functions.

In this manuscript, we show that vRNP/Rab11 hotspots constitute viral inclusions that are not delimited by membranes and display characteristics of liquid organelles. Liquid properties include dynamic change of components, round appearance, easy deformation upon application of sheer force or fusion events and fast adaptation to physiological changes. We show that the liquid organelles are formed in the vicinity of the Endoplasmic Reticulum Exit Sites (ERES) (or transitional ER) and their assembly is dependent on continuous ER-Golgi vesicular cycling. We demonstrate that, contrary to the current view, these sites are not formed by established RNA-RNA interactions amongst different vRNP segments, but precede viral genome assembly. We propose that the condensed IAV inclusions do not promote escape to the antiviral response but facilitate stochastic RNA-RNA interactions in a liquid environment of crowded vRNPs.

## Results

### vRNPs and Rab11 form rounded viral inclusions that are not membrane delimited

Using electron microscopy, we previously showed that viral infection induced clustering of vesicles heterogeneous in size^14^. These constitute, in high percentage, round-shaped molecular concentrates, enriched in membranes at the core, but interestingly not delimitated from the cytosol by membranes (Fig. 1a, yellow arrows and quantification in Supplementary Fig. 1). Such structures are found in cells infected by many viruses and known as viral inclusions, as they concentrate viral (and cellular) material^24,25^. In agreement, using correlative light and electron microscopy, areas of clustered vesicles/viral inclusions matched those of vRNPs and Rab11 identified by immunofluorescence^14^. Using double immunogold labelling, we confirmed the existence of electron dense regions (green arrowheads) positive for vRNPs, protruding from vesicles (red arrowheads) positive for Rab11 (Fig. 1b). To investigate whether the lack of membrane enabled viral inclusions to react fast to physiological changes or whether they constituted crystallized aggregated material, we subjected them to hypotonic shock. Infected cells expressing GFP-NP to label vRNPs^30^ were live-imaged by confocal microscopy. After approximately 1 min, cells were subjected to a rapid hypotonic shock (the ionic strength changed from 150 to 0.300 mM by diluting media with water). Viral inclusions, otherwise stable over time, immediately started to dissolve, and 2 min later were no longer visible (Fig. 1c, Supplementary Movies 1 and 2). The ability of the IAV viral inclusions to react to dilution suggests a liquid character^31^. Together, these data reveal that viral inclusions, containing both Rab11 and vRNPs, can respond to changes in the cellular environment and their constituents can self-organize into fluxional spherical structures in live cells, behaving like a membraneless organelle.

**Figure 1.**
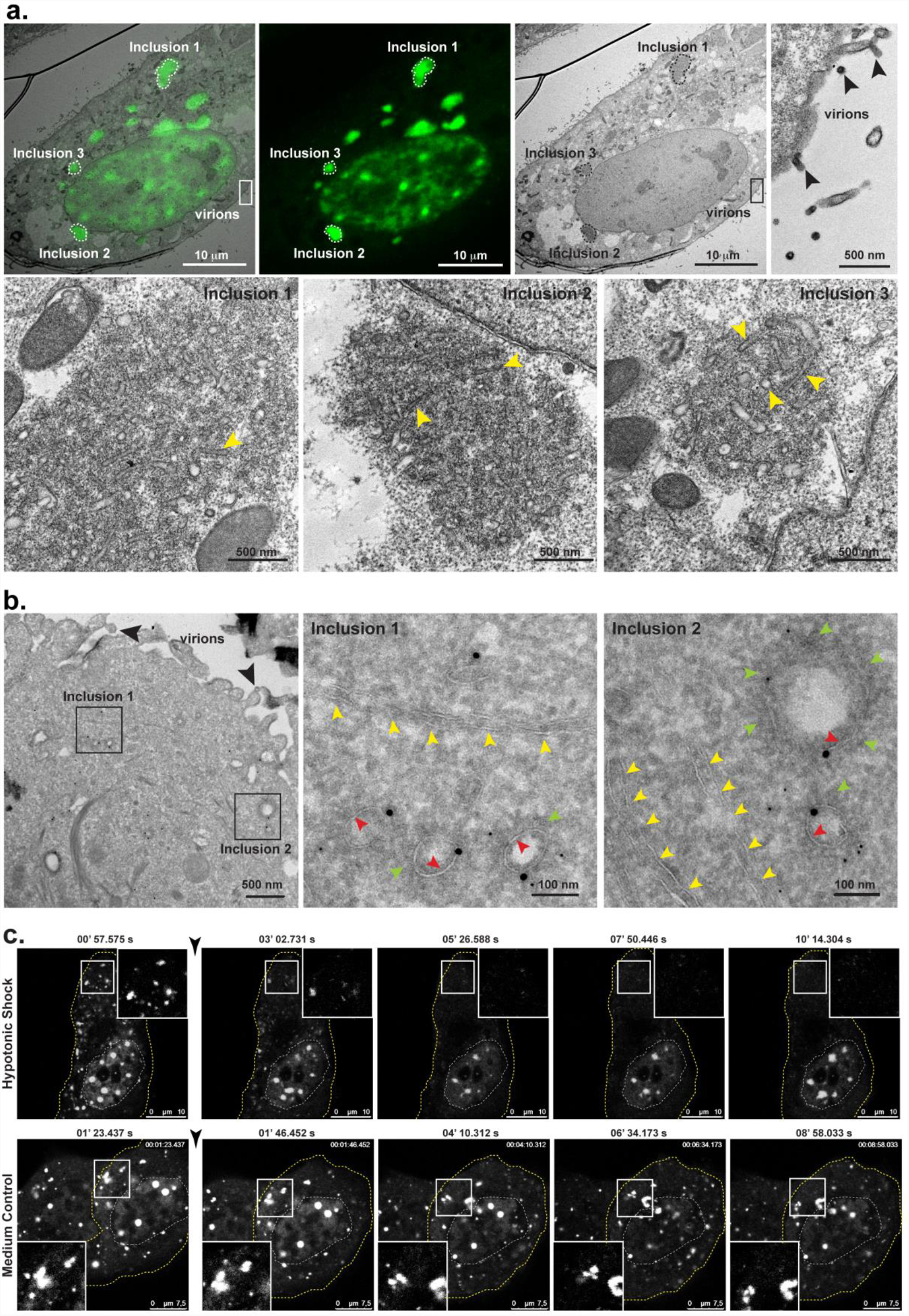
vRNPs and Rab11 form membraneless organelles that quickly respond to changes in the cellular environment. **a.** HeLa cells were transfected with a plasmid encoding GFP-NP and co-infected with PR8 virus, at an MOI of 10. Cells were imaged by confocal and electron microscopy and the resultant images were superimposed. Areas of correlation, inclusions 1 to 3, are delineated by a dashed line in the upper panel and shown in greater detail in the lower panel. Progeny virions budding at the surface (black arrowheads) show that the cell was infected. Yellow arrowheads show individual vesicles within the inclusion. Bar = 10 μm or 500 nm. **b.** GFP-Rab11 WT cells were infected with PR8 virus, at an MOI of 5, for 16 h. Cells were stained for GFP (18 nm gold particles) and viral NP (6 nm gold particles). Inclusion areas are highlighted by black boxes. Yellow arrowheads indicate ER structures in the vicinity of viral inclusions. Black arrowheads show progeny virions budding at the cell surface. Red arrowheads show Rab11 vesicles. Green arrowheads show electron-dense vRNPs. Bar = 100 or 500 nm. **c.** A549 cells were transfected with a plasmid encoding GFP-NP and co-infected with PR8 virus, at an MOI of 5. At 16 hpi, cells were imaged under time-lapse conditions. The black arrowhead indicates addition of water (hypotonic shock) or regular growth medium. White boxes highlight vesicular clusters in the cytoplasm in the individual frames. The dashed white line marks the cell nucleus, whereas the dashed yellow line delineates the cell periphery. Bar = 7.5 or 10 μm. Images were extracted from Supplementary Movies 1 and 2. Experiments were performed at least twice.

### vRNPs form viral inclusions with properties of liquid organelles

Many membraneless organelles have been described in the cell, being supramolecular assemblies formed by nucleic acids and proteins^32^. Examples include nuclear speckles, nucleolus, centrioles and stress granules^33,34^. Interestingly, these were shown to have liquid-like properties on account of their dynamic exchange of material, ability to internally reorganize, rounded shape, and deformability promoted by fusion and fission events^32,35^. Recently, Negri bodies formed during rabies virus infection were shown to have liquid-like properties^27^, and it was postulated that other viral factories or viral inclusions among *Mononegavirales* would assemble by liquid-liquid phase separation. To test this idea for influenza A viral inclusions, we analysed their dynamic nature inside living cells. We observed that viral inclusions were highly dynamic, as demonstrated in Fig. 2a (and Supplementary Movie 3). To capture the movement of a 1 min movie in a snapshot, we show the average intensity of labelled vRNPs (in red) as a defined puncta, surrounded by a wider green area that corresponds to the standard deviation of the average, indicating that these structures are highly dynamic (Supplementary Movie 3). We then enquired if individual clusters exchanged material with the exterior and performed fluorescence recovery after photobleaching (FRAP). We found a high variation in the behavior of cytosolic viral inclusions, with different speeds and patterns of recovery of the fluorescent signal. Some exhibited a fast and complete recovery (Fig. 2b, purple line, Supplementary Movie 3), whilst others showed a slower and incomplete recovery (Fig. 2b, blue line, Supplementary Movie 3). The recovery profile was also variable with some regions losing and/or gaining intensity during the recovery phase (Fig. 2b, purple line) but others exhibiting a steady progression of fluorescence recovery (Fig. 2b, blue line). Not surprisingly, when the collection of FRAP events was averaged, the recovery profile obtained had a very large standard deviation (Fig. 2c). The calculated half time of recovery was 2.9 seconds and the diffusion rate calculated was 2.422 ± 0.154 m^-13^ s^-1^ (D ± SEM), a value similar to what has been found for other liquid organelles including Negri bodies formed during rabies virus infection^27^. These measurements are also consistent with those of nucleoli and stress granules^27,36^, indicating that viral inclusions exchange material with similar structures or with the cytosol. The mobile fraction of vRNPs varied from 41.4 ± 15.7% (mean ± SD) at 5 seconds to 63.5 ± 39.7% within 60 seconds, and the curve plateaued after 15 seconds (Fig. 2c). The immobile GFP-NP must be biologically relevant. It either translates thermodynamically stabilized or kinetically trapped state of the protein, as in stable interactions (presumably among different vRNPs in each cluster), or a complex pattern of exchange between vRNPs and the exterior. In fact, careful analysis of dynamic events of individual viral inclusions revealed a constant flux of small material in and out of these rounded structures (Fig. 2d, arrows, Supplementary Movie 4), many fusion events amongst individual inclusions either separated at short or long distance (Fig. 2e, Supplementary Movies 5 and 6) and fission events (Fig 2f, Supplementary Movie 6). Upon fusion/fission events, the rounded shape was reacquired, indicating occurrence of internal rearrangements. Some acquisition of material originated from several compartments being difficult to track their origin (Fig. 2e, complex, Supplementary Movie 6). Collectively, these data suggest that vRNP/Rab11 inclusions are liquid droplets arising from phase separation. Furthermore, these inclusions appear to be constantly exchanging material amongst them, which would be essential if these were sites devoted to viral genome assembly.

**Figure 2.**
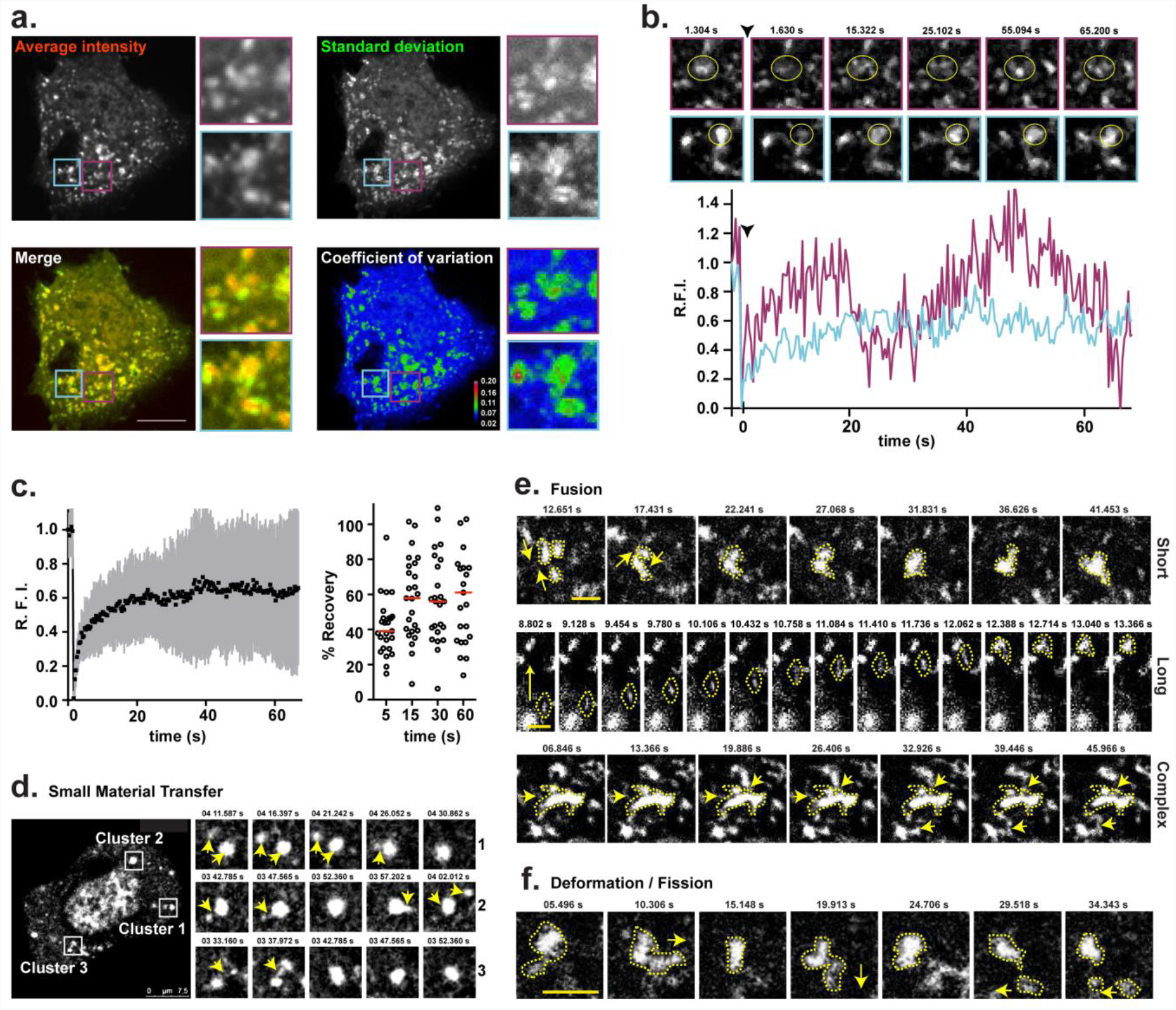
vRNP/Rab11 inclusions have properties of liquid organelles and exchange material dynamically. A549 cells were transfected with a plasmid encoding GFP-NP and co-infected with PR8 virus, at an MOI of 5, for 16 h. Cells were imaged under time-lapse conditions. **a.** A representative infected cell is shown. The fluorescence signal of viral clusters in this cell is depicted as: average intensity (in red), standard deviation (in green), the merge of both, and coefficient of variation. Two areas of viral NP inclusions, highlighted in purple and cyan boxes, were selected for fluorescence recovery after photobleaching (FRAP). Bar = 10 μm **b.** The photobleached regions are marked by a yellow circle. The black arrowhead indicates the time of photobleaching. Relative fluorescence intensity (R.F.I.) was plotted as a function of time for each particle. Images have been extracted from Supplementary Movie 3. **c.** R.F.I. was plotted as a function of time for the means of 25 FRAP events (left graph). The means are shown (black) with error bars representing the standard deviation (gray). The percentage of recovery of each photobleached region is shown for specific times (right graph), with means represented as red bars. A single experiment representative of two independent experiments is **d.** A representative infected cell is shown in the large image, with selected clusters marked by white boxes. Individual frames with single moving particles, from each cluster, highlighted with yellow arrows are shown in the small panels. Bar = 7.5 μm. Images were extracted from Supplementary Movie 4. **e.** Individual frames show three distinct fusion events: at short distance – simple and complex – and at long distance. **f.** Similarly, deformation and/or fission event is shown. Yellow arrows highlight fusion (**e.**) or fission (**f.**) movements, whereas yellow dashed lines indicate the shape of clusters. Images were extracted from Supplementary Movies 5 and 6. Bar = 2 μm.

### IAV viral inclusions form when a single vRNP is expressed in cells

The most accepted model establishes that RNA-RNA interactions lead to the formation of viral inclusions. However, the dynamic exchange of material within clusters supports that these structures promote genome assembly. The formation of viral inclusions, measured by redistribution of Rab11, was reported on a mini-replicon system expressing only two vRNPs, segment 7 and 8^10^, for which no particularly strong RNA-RNA interactions have been demonstrated^37,38^. In addition, vRNPs self-repulse and compete to avoid repetition of a vRNP in a virion^39-41^. We tested whether RNA-RNA interactions were needed to form the liquid viral inclusions by assessing the formation of vRNP hotspots and the sub-cellular distribution of Rab11 in the same mini-replicon system expressing one or two segments. Cells were transfected with plasmids expressing the RdRp and NP (3P-NP), NS2 (to ensure nuclear export of vRNPs), segment 7 (that encodes for M1 and M2) and, when indicated, segment 8 (that expresses NS1 and NS2). Segment transcription originates a complete negative sense RNA, to which the RdRp binds, amplifying the system, mimicking viral transcription and replication. As control, the same system without the polymerase PB2 was evaluated (2P-NP).

Results show that Rab11 subcellular distribution did not change in any of the 2P-NP conditions, consistently with previous reports^10^. However, in the 3P-NP condition, Rab11 redistributed, forming the characteristic enlarged puncta regardless of expressing one or two vRNPs, indicating that one vRNP is sufficient to form viral inclusions (Fig. 3a). The increase in areas of Rab11 puncta was significantly different between the 3P-NP and 2P-NP conditions when quantified and ranked based on their size: small inclusions up to 0.15 μm^2^, intermediate inclusions between 0.15 and 0.30 μm^2^, and large inclusions bigger than 0.30 μm^2^ (Fig. 3b), as before^14^. Consistent with our own work, if vRNPs did not exit the nucleus by not including NS2, Rab11 distribution was similar to the 2P-NP condition (Fig 3b, 3P-NP seg7 without NS2). Similarly, in the case of 3P-NP, and independently of the number of segments expressed, vRNPs were detected in puncta, rather than dispersed, showing that vRNP hotspots are formed without requiring RNA interactions among distinct segments (Fig. 3c, upper panels). In the case of 2P-NP conditions, probes against the vRNA of segments 7 or 8 detected discrete dots in the nucleus (Fig. 3b, lower panels), consistent with pol I transcription and lack of amplification, as described before^10^. The expression of all components of each condition was evaluated by western blotting, except that of NS2, for which no good commercial antibody is available (Fig. 3d). Confirming that the system was fully functional, the corresponding proteins of a specific segment were detected only in 3P-NP samples (Fig. 3d).

**Figure 3.**
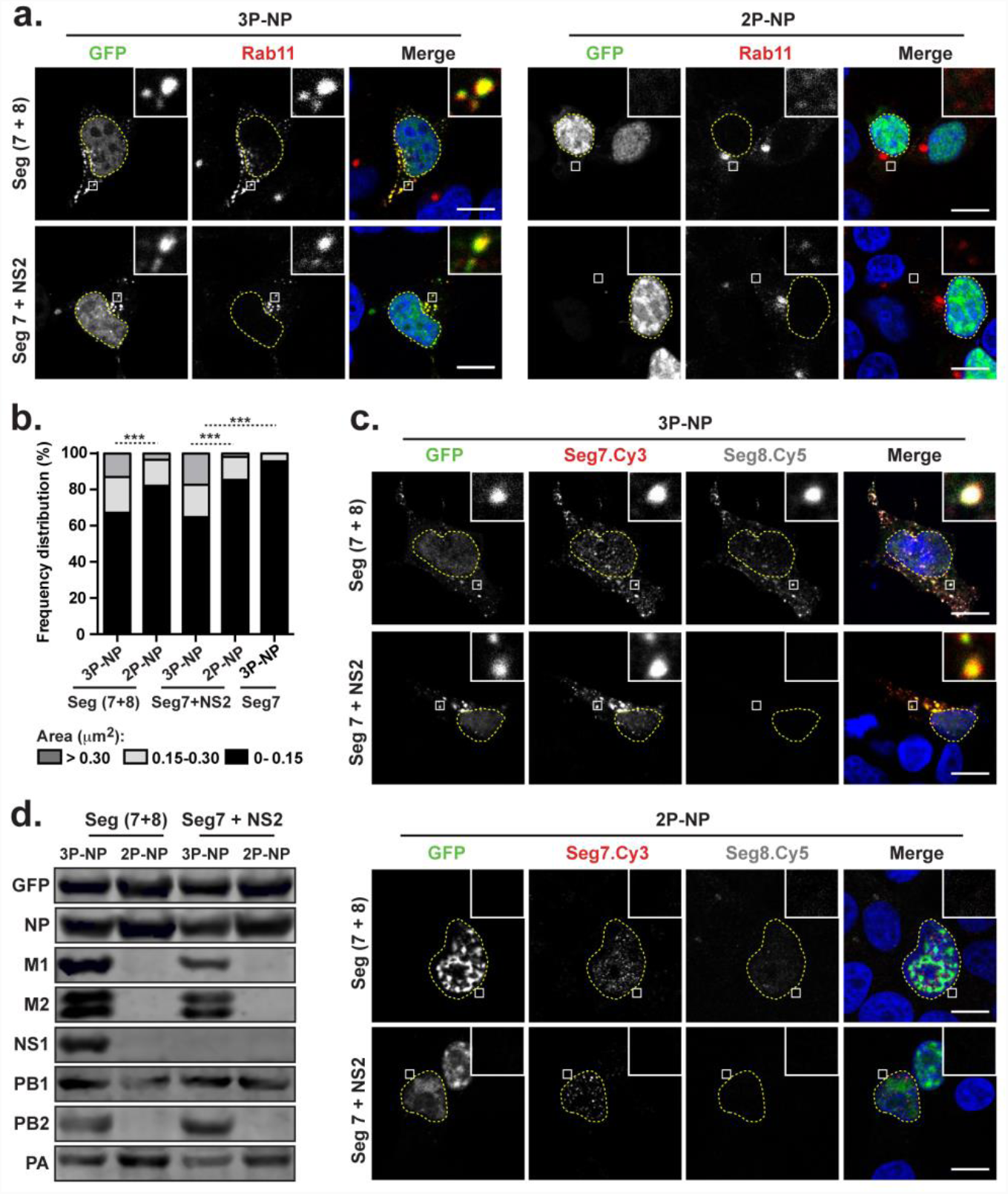
Viral inclusions form in the absence of RNA-RNA interaction. 293T cells were transfected for 16 h with plasmids expressing vRNA segments 7 and 8, or segment 7 alone, and the minimal protein components of an influenza RNP: the three polymerase proteins (3P) (or, as a nonfunctional control, two polymerase proteins lacking PB2 - 2P) and NP, as well as with plasmids expressing GFP-NP. Cells were also transfected with a plasmid encoding NS2, when segment 7 was expressed alone. **a.** Cells were fixed and stained for Rab11 (red). White boxes show areas of co-localization between NP and Rab11. Nuclei are delineated by yellow dashed lines. Bar = 10 μm. **b.** The frequency distribution of Rab11 inclusions within the three area categories (in μm^2^) was plotted for each condition. Statistical analysis of data was performed using a non-parametric Kruskal– Wallis test, followed by Dunn’ s multiple comparisons test (***p < 0.001). Statistical analysis compares the area of all inclusions between conditions. Between 30 and 70 cells were analyzed per condition. **c.** Duplicate samples were processed to detect segment 7 (red) and segment 8 (gray) RNA by FISH. White boxes show areas of co-localization between NP and viral segments. Nuclei are delineated by yellow dashed lines. Bar = 10 μm. **d.** Cells were lysed and indicated proteins were detected by Western blotting.

Collectively, the obtained results demonstrate that viral inclusions assemble in the presence of a single vRNP. The data indicates that formation of Rab11 enlarged puncta is dependent of vRNPs reaching the cytosol, but precedes and is not dependent on RNA-RNA interactions.

### vRNP-containing viral inclusions are not responsible for escaping innate immunity activation

It has been shown that phase-separated compartments are able to sequester or exclude specific material, including components of the innate antiviral immune response^42,43^. It is therefore possible that formation of IAV viral inclusions is a strategy to prevent the activation of cell-intrinsic defenses, either by sterically excluding sensors of exogenous material or by sequestering key factors of the downstream pathways. To address this hypothesis, we have used A549 cells constitutively expressing a fully functional or a non-functional form of GFP-tagged Rab11 (GFP-Rab11 wild type (WT) or GFP-Rab11 dominant negative (DN), respectively). These cells were established in our lab, and have been evaluated for growth rate and permissiveness to viral infection^14^. Both cell lines were infected with WT PR8 or an NS1 mutant virus that does not express a functional form of the main viral factor supressing cell antiviral responses (NS1-N81)^44^. To characterize viral infection, cells were fixed at 8 and 16 h post-infection (hpi), stained for NP protein and imaged by confocal microscopy (Fig. 4a). Changes in Rab11 subcellular distribution were quantified by measuring the area of Rab11 inclusions in infected and control cells, and ranking them as above (Fig. 4b). As previously published by us^14^, infection of cells stably expressing GFP-Rab11 WT with WT PR8 virus induced a redistribution of Rab11, forming viral large inclusions that contained vRNPs (Fig. 4b). Furthermore, the frequency of large Rab11 inclusions increased as infection progressed from 8 to 16 hpi. Noteworthy, infection of this cell line with the NS1 mutant virus produced similar changes in the frequency distribution of the different size category inclusions (Fig. 4b). Infection of GFP-Rab11 DN cell line, either with WT PR8 or NS1 mutant virus, did not change Rab11 DN distribution (Fig. 4a,b). Consistent with previous reports^12,17,45^, Rab11 DN was primarily localized to the TGN, with some diffuse cytoplasmic staining also visible (Fig. 4a). Also in agreement with these studies, overexpression of Rab11 DN impaired the formation of viral inclusions characteristic of IAV infection and formation of vRNP hotspots^11,13^, and therefore NP was diffusely distributed throughout the cytoplasm (Fig. 4a). Next, we examined the impact of constitutively expressing GFP-Rab11 WT or DN on viral replication, by plaque assay (Fig. 4c). WT PR8 and NS1-N81 virus production in GFP-Rab11 DN cell line was significantly impaired when compared with GFP-Rab11 WT cell line, with an approximately 50% reduction in viral titres at 16 hpi (Fig. 4c). These results corroborate previous studies showing that a fully functional Rab11 protein is required for efficient infectious virus production^12,45^. As expected^46^, NS1 mutant virus replication was attenuated in both cell lines, as compared to WT PR8 virus (Fig. 4c). In order to investigate if impaired formation of viral clusters resulted in enhanced activation of the interferon (IFN) cascade, the transcript levels of type I (IFN-α and IFN-β) and type III (IL-29) IFN, and of the IFN-stimulated gene viperin, were analysed at 8 and 16 hpi. For positive control, cells were transduced with the double-stranded RNA mimic polyinosinic:polycytidylic acid [poly(I:C)]. Results show that there are no differences between both cell lines, in any of the conditions analysed (Fig 4d). Also, NS1 mutant virus induced higher mRNA levels than the WT virus, confirming the IFN-antagonizing role of NS1 (Fig 4d). At the protein level, cell lysates from infected cultures were probed for active, phosphorylated IFN regulatory factor 3 (IRF3), a hallmark of activation of the IFN induction cascade (Fig. 4e), and cell culture media were tested for the levels of secreted IFN-β (Fig. 4f). Again, the results obtained were identical for both cell lines. In summary, all results point towards innate immune responses not being affected by biological phase transitions of vRNPs.

**Figure 4.**
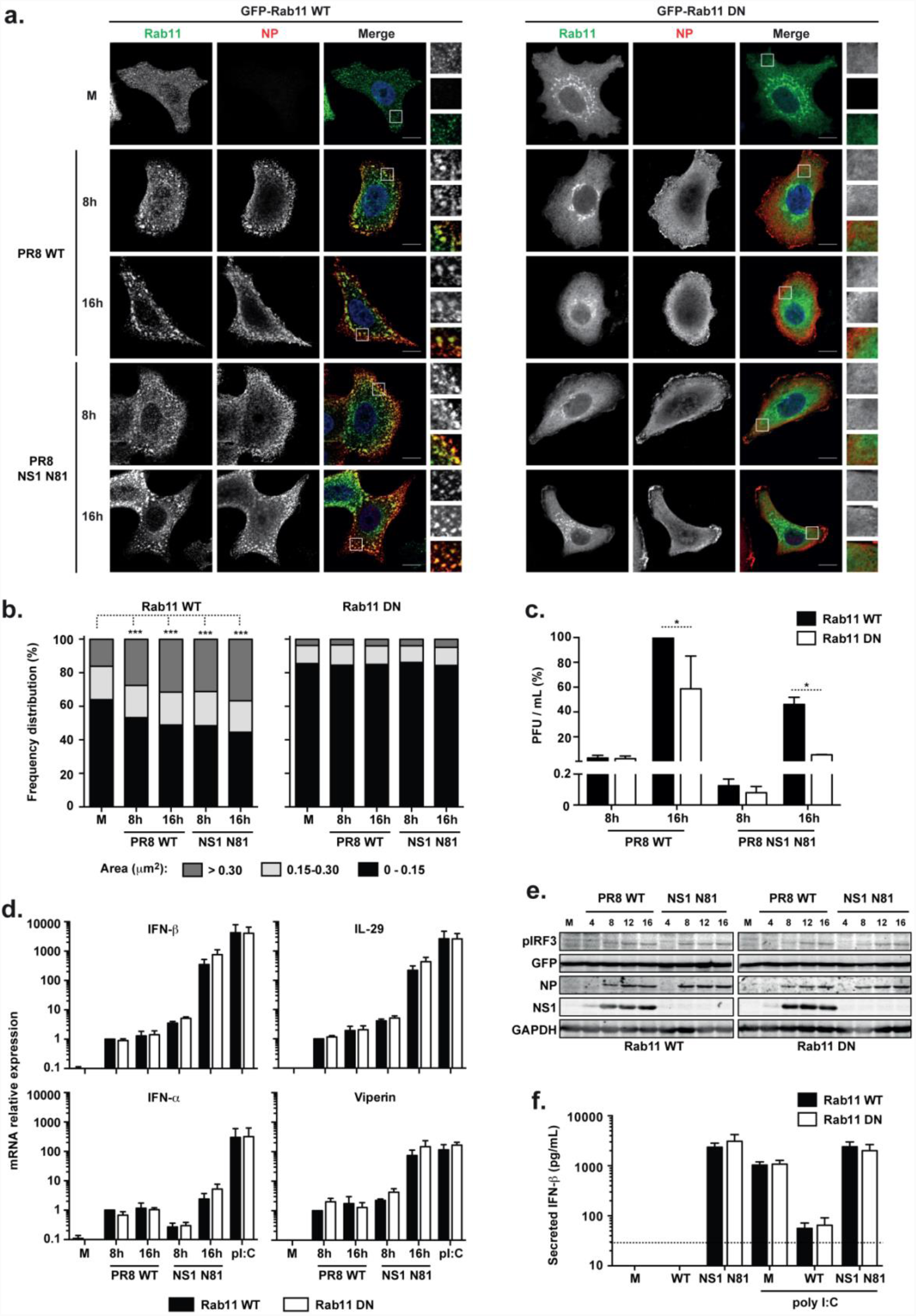
Cell antiviral defenses are not affected by the formation of viral inclusions. **a.** GFP-Rab11 WT and GFP-Rab11 DN cells were infected or mock-infected (M), at an MOI of 3, with PR8 WT or NS1-N81 viruses. Cells were fixed at the indicated times and stained for NP (in red). Bar = 10 μm. **b.** The frequency distribution of NP inclusions within the three area categories (in μm^2^) was plotted for each cell line. Statistical analysis of data was performed using a non-parametric Kruskal–Wallis test, followed by Dunn’ s multiple comparisons test (*** p < 0.001 for GFP-Rab11 WT cells; no statistical significance found for GFP-Rab11 DN cells). Statistical analysis compares the area of all inclusions between conditions. An average of 30 cells was analyzed per condition. A single experiment representative of two independent experiments is shown. **c.** At the indicated times, supernatants were collected and viral production was evaluated by plaque assays using MDCK cells. Statistical analysis of data was performed using two-way ANOVA test, followed by Sidak multiple comparisons test (*p < 0.05 at 16 hpi; no statistical significance found at 8 hpi). Data represents the average of three independent experiments. **d.** Expression of IFN-β, IFN-α, IL-29 and viperin was evaluated at the level of transcription by RT-qPCR in relation to GAPDH. Poly(I:C) was used as a positive control for maximum expression of these transcripts. Statistical analysis of data was performed using two-way ANOVA test, followed by Sidak multiple comparisons test (no statistical significance between conditions found). Data represents the average of three independent experiments. **e.** Expression of phosphorylated IRF, GFP, NP, NS1 and GAPDH was evaluated at the protein level by western blotting. **f.** The levels of secreted IFN-β were quantified by ELISA in cell supernatants at 24 hpi. Poly(I:C) was used as a positive control for maximum expression of IFN-β protein. The limit of detection of this method is 30 pg/mL (dashed line). Statistical analysis of data was performed using two-way ANOVA test, followed by Sidak multiple comparisons test (no statistical significance between conditions found). Data represents the average of three independent experiments.

### Viral inclusions form in the proximity of the ER, displaying matching movements

Given that phase separation is generally a spatially regulated process, we next asked whether this was the case for IAV viral inclusion assembly. It has recently been reported that vRNPs associate with the ER when leaving the nucleus and that Rab11 would collect vRNPs from the ER for delivery to the surface^16^. Our electron microscopy data also indicates that the ER is constantly found in close proximity to viral inclusions (Fig. 1b, yellow arrowheads). We therefore tested if clusters are associated with the ER, by using antibodies against different ER markers or a cell line expressing a fluorescent tagged-ER membrane marker (HeLa Sec61β-Emerald)^47^. Confocal imaging of cells at different times post-infection failed to identify co-localization between the ER and viral inclusions (Supplementary Fig. 2a-c). However, it was evident that, from 8 hpi onwards, the vRNP clusters dispersed throughout the cytoplasm were frequently found juxtaposed to ER tubules (Supplementary Fig 2a-c, inlets), suggesting an association between both structures. To gain insight into the dynamics of ER-viral inclusion association, live cell imaging was performed. For this, HeLa Sec61β-Emerald cells were transfected with mCherry-NP and infected with PR8 (Fig. 5a and Supplementary Movie 7), or A549 cells were co-transfected with mCherry-NP and ER-GFP and infected with PR8 virus (Fig. 5b and Supplementary Movie 8). In both experiments, viral inclusions displayed movements that matched those of the ER, although it is not clear whether ER motion was driving displacement of viral inclusions or, conversely, viral inclusions were gliding over the surface of the ER tubules. The ER is a complex organelle, with distinct morphologies and diverse functions^47-49^. In order to identify the specific ER domain interacting with viral inclusions, we tested different markers, including Atlastin 3, which accumulates in 3-way junctions, Sec23 and Sec31, both present in ER Exit Sites (ERES). We observed no correlation between Atlastin 3 and vRNPs staining (Supplementary Fig. 2d), but Sec23 and Sec31 localized between the clusters and the ER, and even co-localized with NP in specific spots (Fig. 5c, Supplementary Fig. 2e). Live cell imaging of Sec16, another ERES component, indicates that these structures serve as docking platforms where vRNPs accumulate, allowing a constant flux of material in and out of viral inclusions, with frequent fission and fusion events taking place (Fig. 5d, Supplementary Movies 9 and 10).

**Figure 5.**
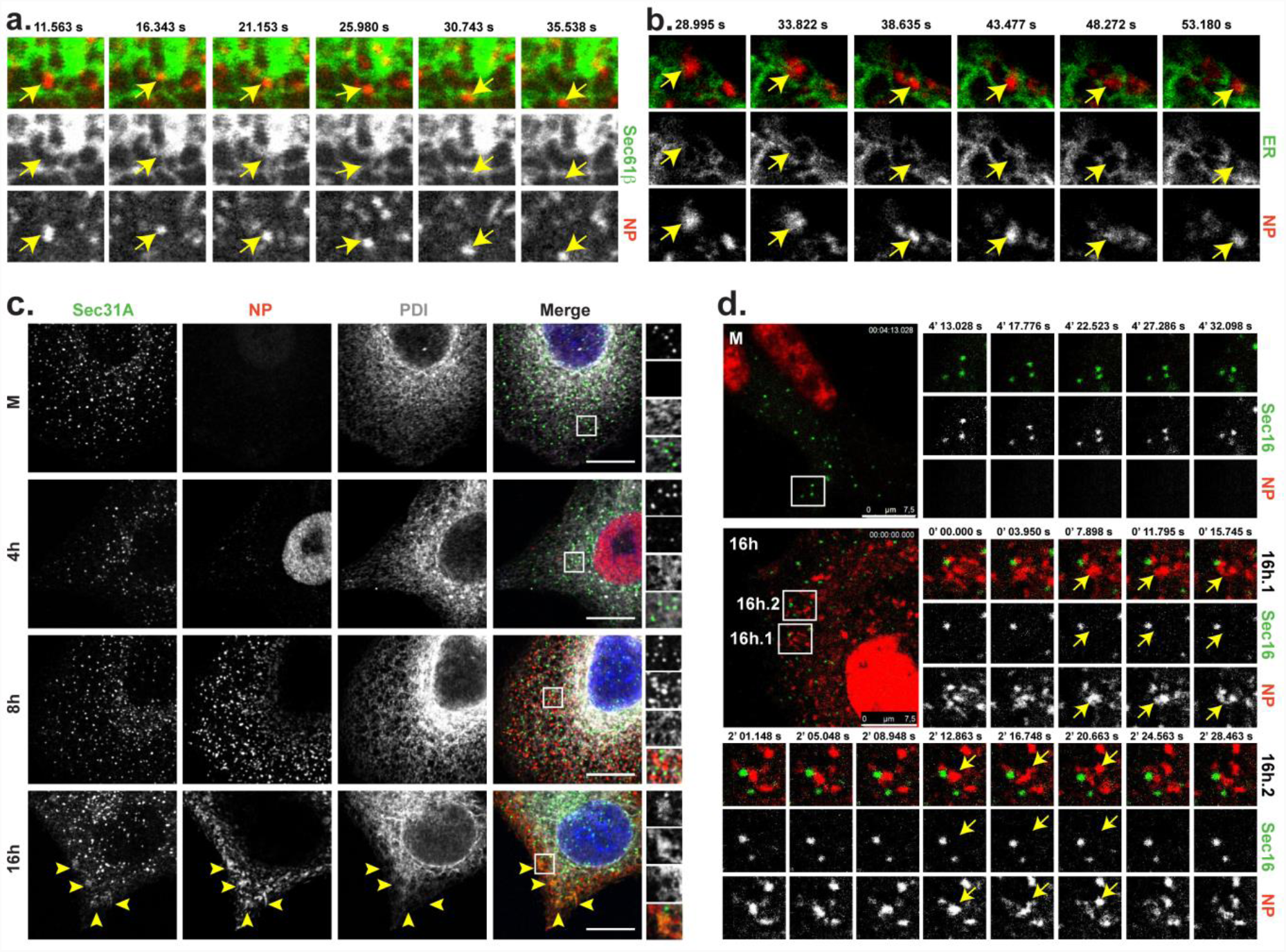
Viral inclusions are associated with ER exit sites. **a.** Sec61β-Emerald cells were transfected with mCherry-NP and infected with PR8 virus, at an MOI of 10, for 16 h. **b.** A549 cells were co-transfected with plasmids encoding mCherry-NP and ER-GFP and infected with PR8 virus, at MOI of 10, for 16h. **a., b.** Cells were imaged under time-lapse conditions. Individual frames with single moving particles highlighted with yellow arrows are shown in the small panels. Bar = 2.5 μm. Images were extracted from Supplementary Movies 7 and 8. **c.** A549 cells were infected or mock-infected (M) with PR8 virus, at an MOI of 3, and fixed at the indicated times. Cells were stained for the ER proteins Sec31 (in green) and PDI (in gray) and the viral NP protein (in red). Areas highlighted by the white box are shown on the right of each panel. Bar = 10 μm. **d.** A549 cells were co-transfected with plasmids encoding mCherry-NP and GFP-Sec16 and infected or mock-infected (M) with PR8 virus for 16 h. Cells were imaged under time-lapse conditions. Representative cells are shown in the left large images. Individual frames with single moving particles highlighted with yellow arrows are shown in the small panels. Two examples are provided for the infected cell (16h.1 and 16h.2). Bar = 7.5 μm. Images were extracted from Supplementary Movies 9 and 10.

### Continuous Golgi-ER vesicular cycling controls the formation of viral inclusions

The ERES are specialized domains where secretory proteins are loaded into coat protein complex II (COPII)-coated vesicles and transported to the Golgi^50^. Recruitment of COPII proteins to the ERES is controlled by the Sar1 GTPase cycle^51^. This small-GTPase also regulates ER-membrane tubulation and vesicle fission, having a critical role in the generation of the ERES^52^. To analyse the effect of disrupting the ERES on the assembly of viral inclusions, we overexpressed a GTP-restricted mutant of GFP-tagged Sar1 (Sar1-GTP), which inhibits anterograde protein transport. Overexpression of GFP and GFP-tagged Sar1 WT were performed as controls. Immunofluorescence analysis showed that overexpression of Sar1-GTP strongly reduced the size of viral inclusions, when compared to overexpression of GFP or Sar1, in a statistically significant manner (Fig. 6a, b). The number of inclusions per μm^2^ of cellular area and the percentage of NP signal that is inside viral inclusions were also analyzed, with the latter being significantly reduced when Sar1-GTP was overexpressed [7.7 ± 2.9% (mean ± SD) in Sar1-GTP vs 12.7 ± 3.8% in GFP (p<0.0001) and 11.1 ± 3.4% in SAR1 (p=0.0005)] (Fig. 6b). Confocal imaging of the viral transmembrane protein hemaglutinin (HA) confirmed that Sar1-GTP is disrupting ER-Golgi trafficking, since this protein was retained and accumulated in the ER when Sar1-GTP was overexpressed, but reached the plasma membrane during GFP and Sar1 overexpression (Fig. 6c).

**Figure 6.**
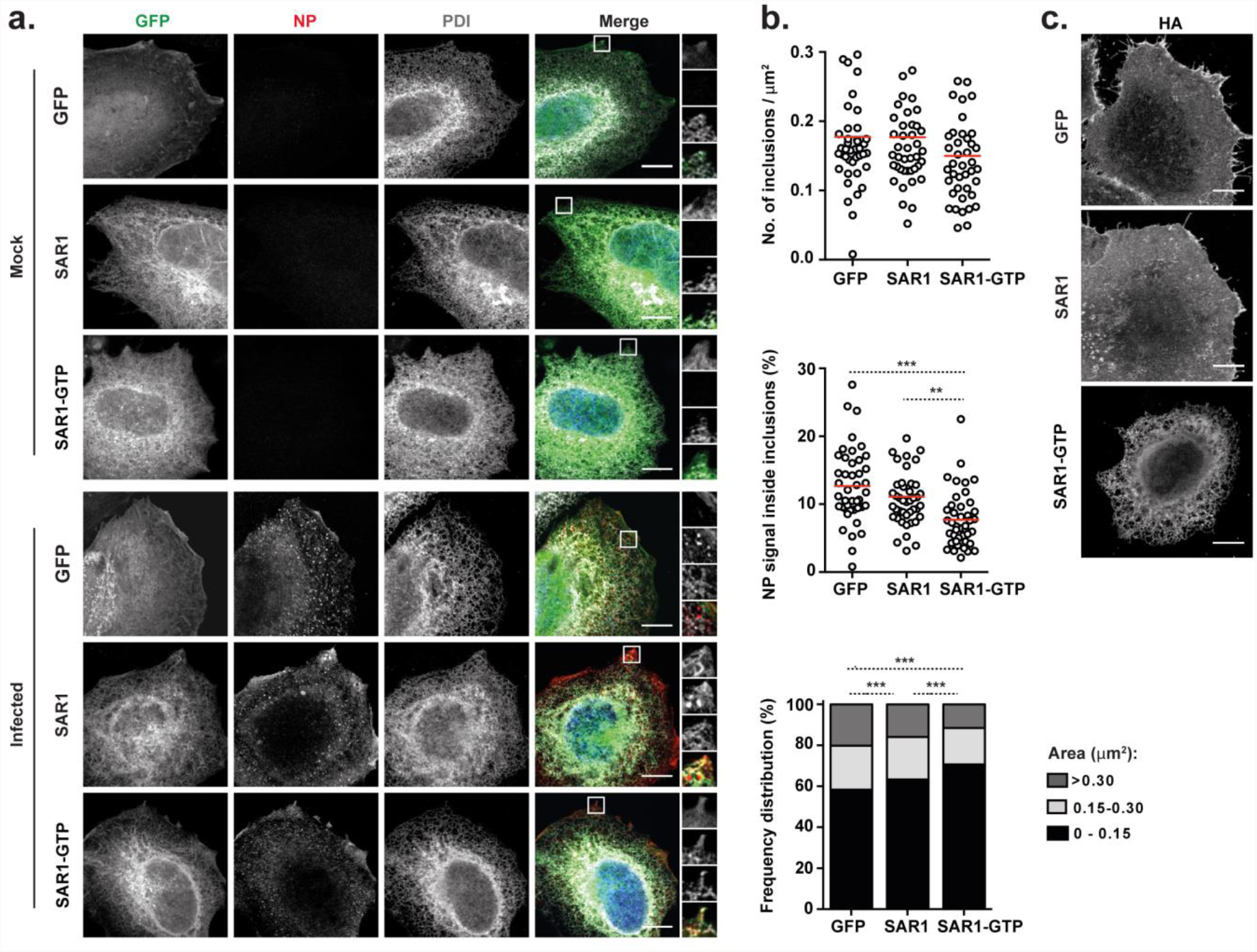
vRNP clustering is impaired by disruption of ER-exit sites. HeLa cells were transfected with plasmids encoding GFP, SAR1 WT-GFP or SAR1 GTP-GFP and, 24 h later, infected or mock-infected with PR8 virus, at an MOI of 10. At 16 hpi, cells were fixed and processed for immunofluorescence. **a.** Cells were stained for the viral protein NP (in red) and for the ER protein PDI (in gray). Areas highlighted by the white box are shown on the right of each panel. **b.** The frequency distribution of NP inclusions within the three area categories (in μm^2^), the number of inclusions per μm^2^, and the percentage of NP staining that is inside inclusions were plotted for each condition. Statistical analysis of data was performed using a non-parametric Kruskal–Wallis test, followed by Dunn’ s multiple comparisons test (***p <0.001). More than 40 cells from 2 independent experiments were analyzed per condition. **c.** Infected cells were stained for the viral protein HA and imaged by confocal microscopy. Bar = 10 μm.

The above results suggest that the establishment of viral inclusions may also require the Golgi compartment. To address this issue, we inhibited the shuttling of cargo proteins between the Golgi and the ER by treating cells with brefeldin A (BFA)^53,54^. Upon addition of a low dosage of BFA to cells infected for 8 h, viral inclusions disassembled within less than 1 hour (Fig. 7a, lower panel). There was a robust and statistically significant decrease in the size [from 0.242 ± 0.203 μm^2^ (mean ± SD) to 0.189 ± 0.154 μm^2^, p<0.0001] and number of viral inclusions per μm^2^ (from 0.264 ± 0.051 to 0.179 ± 0.056, p<0.0001), as well as in the percentage of NP that was inside the inclusions (from 17.5 ± 4.8 to 9.2 ± 2.8%, p<0.0001) (Fig. 7b). Immunostaining of cells with ER and Golgi makers (PDI and GM310, respectively) revealed that BFA treatment provoked the disassembly of the Golgi complex, as expected^53,54^, but not of the ER (Fig. 7a). Areas stained for Rab11 also decreased with BFA treatment (Fig. 7c). Note that the viral transmembrane protein M2 still localized at the plasma membrane, likely because of low dosage (2 μg/mL) and short duration (1 h) of the BFA treatment (Fig. 7c)^53,54^. In sum, the data collectively shows that biogenesis of IAV liquid viral inclusions enriched in vRNPs and Rab11 is dependent on continuous cycles of material between the ER and the Golgi, indicating that its distribution is spatially regulated.

**Figure 7.**
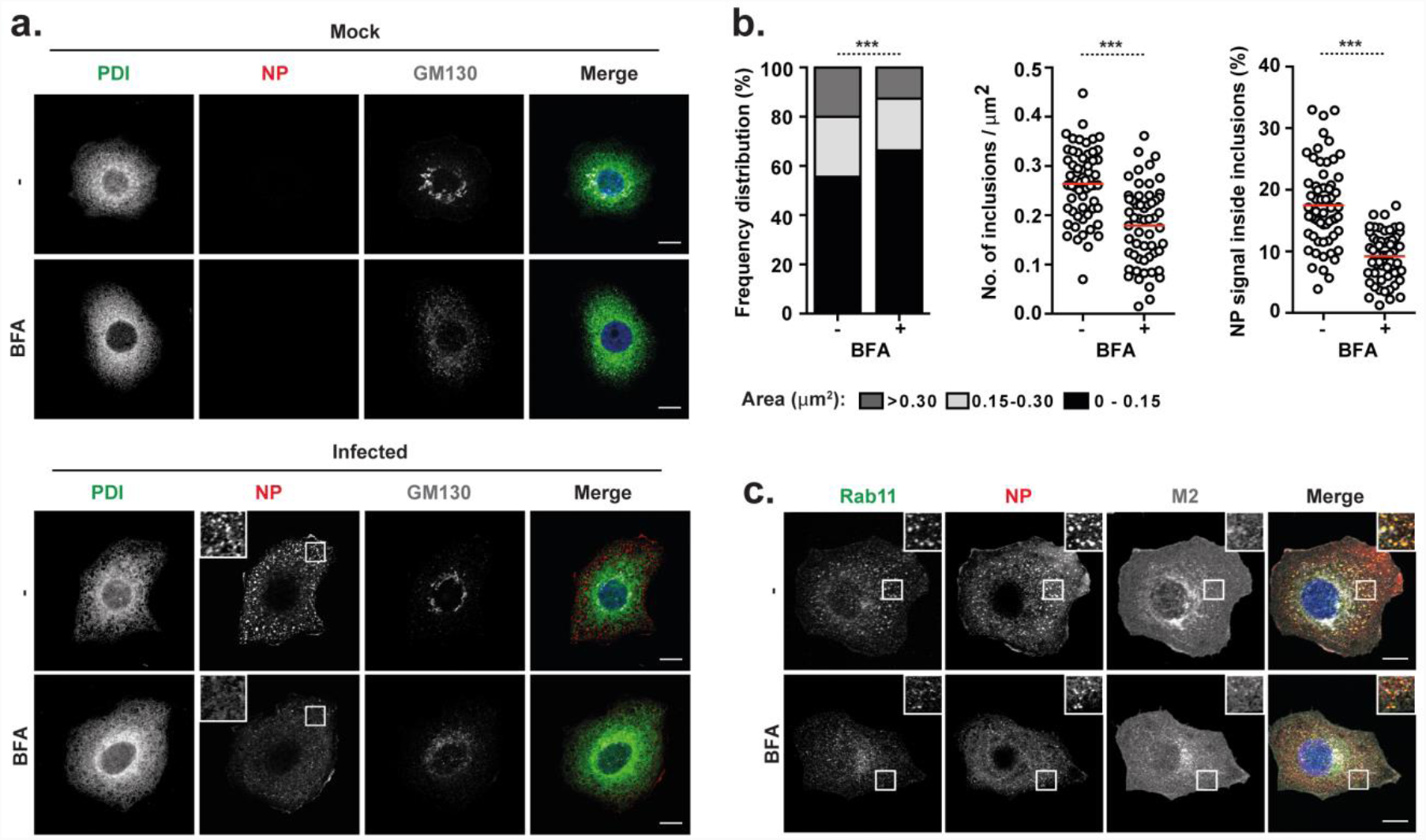
Disruption of ER-Golgi trafficking disassembles vRNP hotspots. A549 cells were infected or mock-infected with PR8 virus at an MOI of 3 for 8 h, and then treated or mock-treated with 2 μg/mL of brefeldin A (BFA) for 1h. **a.** Cells were immunostained for the ER marker PDI (in green), the viral protein NP (in red) and the cis-Golgi marker GM130 (in gray) and imaged by confocal microscopy. Selected areas of the cytoplasm are marked by white boxes and displayed on the top left corner of the images. Bar = 10 μm. **b.** The frequency distribution of NP inclusions within the three area categories (in μm^2^), the number of inclusions per μm^2^, and the percentage of NP staining that is inside inclusions were plotted for each condition. Statistical analysis of data was performed using a non-parametric Kruskal–Wallis test, followed by Dunn’ s multiple comparisons test (***p <0.001). An average of 60 cells from 2 independent experiments was analyzed per condition. **c**. Infected cells were stained for the host protein Rab11 (in green) and the viral proteins NP (in red) and M2 (in gray). Cells were imaged by confocal microscopy. Areas highlighted by the white box are shown on the right top corner of each image. Bar = 10 μm.

## Discussion

Phase separation by liquid demixing drives functional compartmentalization in cells. It allows the dynamic isolation of sets of selected molecules^35^ to carry out activities separated from the surroundings without the need of a membrane^55^. Many membraneless organelles, as the nucleolus^33^ and centrosomes^34^, constitutively exist in the cell, while others appear after an insult, as stress granules^36^, DNA repair foci^56^ or G bodies^57^. These organelles are beginning to be accepted as a disseminated biological fast adaptation mechanism to respond to stimuli^36,56-58^. How viruses take advantage of these phenomena is unclear, despite decades have passed since the realization that many infected cells display viral-induced membraneless territories associated with viral replication, viral assembly, and host immune escape^24,28,29,42^.

Here, we report that IAV leads to formation of cytosolic inclusions with liquid-like properties (Figs. 1 and 2). They have similar physical characteristics to those found in rabies virus-infected cells in terms of shape, dynamism and ability to deform^43^, but are not involved in viral replication as it takes place in the host cell nucleus. In addition, these structures also share properties with other reported liquid bodies including reacting fast to physiological changes^31,59^. Formation of these condensates during IAV infection is dependent on Rab11-GTP and on vRNPs. Despite the common characteristics of the molecules involved in IAV inclusion bodies with those described for other membraneless bodies, including multivalency (Rab11)^60,61^, internally disordered regions (NP)^31,62^, nucleic acids (vRNPs)^63^, and oligomerizing RNA binding proteins (NP)^64^, the rules underlying the formation of IAV liquid organelles and their functions are far from understood.

The IAV-induced inclusions were postulated to originate as vRNPs travelling through the cytosol on Rab11 membranes collided, establishing RNA-RNA interactions *in trans*. Inclusions would contain partial to fully assembled genomes^6,11,13,23^. In this work, we provide evidence that viral inclusions are formed with one vRNP (Fig. 3), which indicates that their formation precedes viral assembly. This supports the idea that viral inclusions could operate as dedicated spots in the cytosol to facilitate establishment of RNA-RNA interactions among the eight different segments. This model still accommodates the hypothesis that vesicular collision could drive genome assembly and is consistent with a selective process for assembling the supra-molecular genomic complex. However, as depicted in the model of Figure 8, these interactions would be restricted in space, taking place in viral inclusions rather than the entire cytosol, and a constant exchange of material between different viral inclusions would replenish vRNP stocks and remove fully assembled genomes by an unclear process. Nevertheless, for the exchange of material to occur, viral inclusions need to move.

**Figure 8.**
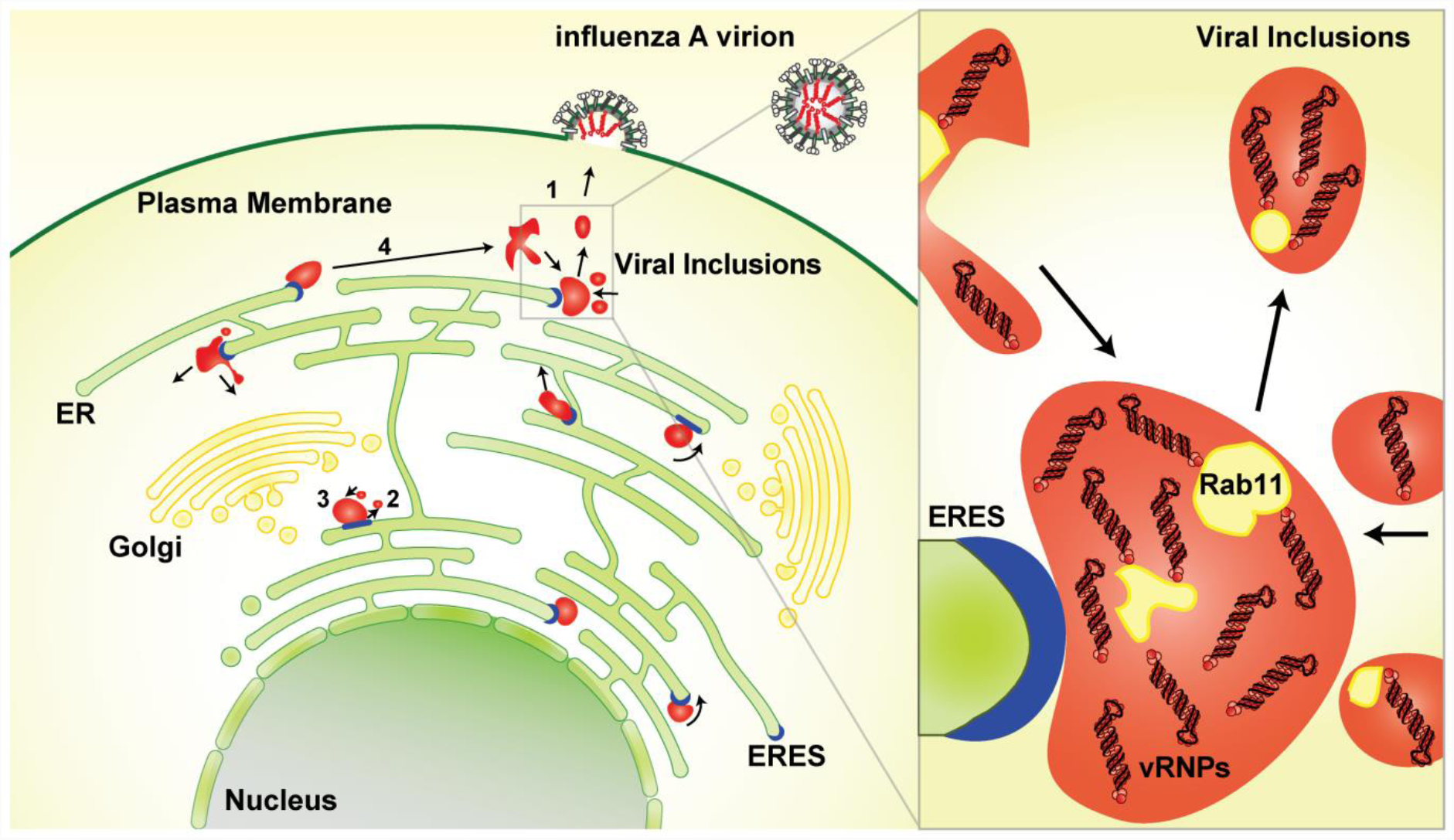
Proposed model. Viral inclusions (in red) exhibit characteristics of liquid organelles, segregating from the cytosol without a delimitating membrane. Viral inclusions exchange material dynamically (1) and deform easily exhibiting fission (2) and fusion (3) events. Viral inclusions can travel long distances before and after fusion/fission events (4), respectively. These organelles are formed in the vicinity of ERES (in blue) and their assembly is dependent on continuous ER-Golgi vesicular cycling. We propose that viral inclusions trigger nucleation of RNA-RNA interactions among the eight different segments to assemble a complete IAV genome. Inlet shows composition of viral inclusions close to ERES. These contain vRNPs of all types, Rab11 and host membranes clustered, but are not delimited by lipid bilayer.

Our work indicates a cross-talk between molecular motors and viral inclusions, with different viral inclusions fusing or dividing in a manner dependent of sheer force (Fig. 2). In addition, IAV Rab11/vRNPs hotspots were shown to exhibit heterogeneous movements, as if moving on actin or sliding on microtubules^10,17,30^, using molecular motors^65^. However, it was recently reported that, in the absence of intact microtubules, vRNAs could be transported through the cytoplasm independently of Rab11-GTP^66^ and a candidate that was recently proposed was the ER^16^. Our work shows, using distinct ways, that molecular crowding occurs in the vicinity of ERES (Fig. 5) and is dependent on continuous vesicular cycling between ER and Golgi (Figs. 6 and 7). This suggests that inclusion formation is spatially regulated. In fact, the movement of IAV inclusions matches that of the ER (Fig. 5) and, in some cases, inclusions seem to slide on ER membranes (Supplementary Movies 7 and 8). This raises the possibility that viral inclusions move gliding through the ER. This notion is in agreement with vRNPs being found at the ER^16^. Interestingly, biogenesis of viral inclusions seems to be intertwined with deregulation of the ERC, which is coherent with a decrease in recycling of transferrin during infection^14,21^, and with Rab11 detection in the ER, as if it was re-directed to this organelle during infection^16^. Whether Rab11 vesicles are targeted to the ER to deliver or to collect vRNPs will be addressed in the future. Inhibiting ER-Golgi vesicle cycles has efficiently locked HA, but not vRNPs, in the ER (Figs. 6 and 7), which supports the former hypothesis, but more experiments are necessary to validate one model. In both cases, there are unresolved questions regarding: 1) the transport of vRNPs to or from the ER; 2) the selection of the ERES as docking sites for viral inclusions and 3) the sensing and transport of fully assembled genomes to the plasma membrane.

Nevertheless, for several biological systems, an intimate association between RNA and liquid membraneless organelles has been reported^57,67-69^. In some cases, RNA promotes phase separation^57,67^, while in other it inhibits this process^68,69^. Many membraneless organelles are involved in RNA metabolism, as is the case of Cajal bodies, nucleoli or stress granules, and therefore it is not surprising that liquid bodies could be involved in IAV genome assembly. Phase separation could however play other roles during IAV infection. We found no differences in the activation of IFN response when viral inclusion assembly was inhibited (Fig. 4), but we have not tested exhaustively other immune related parameters. IAV-induced phase separation leads to localized concentration of vRNPs and many proteins (some possibly unidentified). As shown for other systems, it could operate in signal amplification^70^ or repression/activation of specific cellular pathways by exclusion/inclusion of selected molecules^71^.

Although we favor involvement in viral assembly by spatially restricting vRNPs and facilitate interactions, compelling evidence is required to formally validate this model. Importantly, future experiments will have to detail the internal organization of the viral inclusions. In particular, whether different vRNPs define physical cross-links and establish differences in the property of the material, leading to spatial organization, or alter the diffusion/movement within inclusions, remains to be seen. Besides spatial organization, other factors affecting liquid properties include molecular crowding, solubility affinity or the valency of phase-separating proteins^71^ and these could fluctuate during the course of infection. We are just beginning to understand the involvement of liquid-liquid phase separation in virology, but we anticipate that, given the ancient co-evolution between viruses and eukaryotic cells, and the diversity of host strategies used by viruses, the next years will provide an interesting overlap between the two fields.

## METHODS

### Cells, Viruses and Drugs

The human epithelial cells Madin-Darby Canine Kidney (MDCK), embryonic kidney 293T, cervical HeLa and alveolar basal (A549) were a kind gift of Prof Paul Digard, Roslin Institute, UK. The GFP-Rab11 WT and GFP-Rab11 DN (A549) were produced by our laboratory^14^. The Sec61β-Emerald (HeLa) cell line was a kind gift from Dr Christoph Dehio, Biozentrum, University of Basel, Switzerland^47,72^. All cell types were cultured as described before^10^ and were regularly tested for mycoplasma contamination with the LookOut mycoplasma PCR detection kit (Sigma, MP0035), using JumpStart Taq DNA Polymerase (Sigma, D9307). Reverse-genetics derived A/Puerto Rico/8/34 (PR8 WT; H1N1) was used as a model virus and titrated according to reference^14^. NS1 N81 mutant virus was derived from PR8 WT and expresses only the first 81 amino acids of NS1^44^. Reverse genetic plasmids were contributed by Dr Ron Fouchier, Erasmus MC, Netherlands. Virus infections were performed at a multiplicity of infection (MOI) of 3 to 10. After 45 min, cells were overlaid with DMEM containing 10% fetal bovine serum (Gibco, Life Technologies, 10500-064) and 1% penicillin / streptomycin mix (Biowest, L0022-100). The drug brefeldin A (Sigma) was dissolved in ethanol and used at final concentration of 2 μg/ml.

### Plasmids

HA tagged Sec23 plasmid was a kind gift from Dr Colin Adrain, IGC, Portugal. GFP tagged Sec16 plasmid was purchased from Addgene. GFP-Sec61β was constructed by PCR-amplifying Sec61β from A549 cDNA and cloning it into pEGFP-C2, using *Hin*dIII and *Kpn*I restriction sites. Sar1A was amplified from A549 cDNA and cloned *Xho*I-*Bam*HI restriction sites of pEGFP-N1. GTP-restricted SAR1 (H79G) was produced by site-directed mutagenesis from SAR1 WT-GFP. ER-GFP plasmid was made from pEGFP-C2, by inserting a C-terminal KDEL sequence by site-directed mutagenesis, and an N-terminal ER-signal sequence from calreticulin by oligo-annealing between *Nhe*I and *Age*I restriction sites. Plasmids used for the minireplicon system have been described in reference^10^, except pcDNA3-NS2. The latter was made by PCR amplification of NS2 (from PR8) and insertion into pCDNA3, using *Eco*RI and *Not*I restriction sites. The following primers/oligos were used:

Sec61β Fw: 5’-TAGAAAGCTTCATGCCTGGTCCGACCC-3’

Sec61β Rv: 5’-TCGAGGTACCCTACGAACGAGTGTACTTGCCC-3’

SAR1 WT Fw: 5’-TCGACTCGAGATGTCTTTCATCTTTGAGTGGATCT- 3’

SAR1 WT Rv: 5’- TCGAGGATCCCGGTCAATATACTGGGAGAGCCAGC- 3’

SAR1 H79G FW: 5’- TTTTGATCTTGGTGGGGGCGAGCAAGCACGTCGC - 3’

SAR1 H79G RV: 5’- GCGACGTGCTTGCTCGCCCCCACCAAGATCAAAA - 3’

KDEL Fw: 5’-TGGACGAGCTGTACAAGGACGAGCTGTAATCCGGCCGGACT-3’

KDEL Rv: 5’- AGTCCGGCCGGATTACAGCTCGTCCTTGTACAGCTCGTCCA-3’

Calreticulin tag up: 5’- CTAGCATGCTGCTATCCGTGCCGTTGCTGCTCGGCCTCCTCGGCCTGGCCGTCGCA-3’

Calreticulin tag down: 5’- CCGGTGCGACGGCCAGGCCGAGGAGGCCGAGCAGCAACGGCACGGATAGCAGCATG-3’

NS2 Fw: 5’-CGTAGCGAATTCATGGATCCAAACACTG-3’

NS2 Rv: 5’-GCTAAGACGCGGCCGCTTAAATAAGCTGAAAC-3’

### Transfections

Cells, grown to 70% confluency in 24 well plates, were transfected with 250 ng of indicated plasmids or 100 ng of the synthetic dsRNA polyinosinic:polycytidylic acid [poly(I:C); Calbiochem], using Lipofectamine LTX (Life Technologies) and Opti-MEM (Life Technologies), according to manufacturer’ s instructions. Cells were infected or mock-infected 16 h post-transfection or simultaneously with transfection (live-cell imaging) at indicated MOI.

To reconstitute GFP-tagged RNPs, 293T cells grown to 70% confluency in 24 well plates were transfected with plasmids pcDNA PB1, PB2, PA (130 ng each), NP (150 ng), GFP-NP (50 ng), pPol I segments 7 and 8 (130 ng each) or/and pcDNA-NS2, using Lipofectamine 2000 (Invitrogen) according to the manufacturer’s instructions, incubated overnight, and imaged around 12-16 h later.

### Confocal fixed-cell imaging

Fluorescent *in situ* hybridization (FISH) assay was done as in^10^. Immunofluorescence assays were performed as in^10^. Antibodies used were: rabbit polyclonal against Rab11a (1:100; Life Technologies, 715300), HA tag (1:500; Abcam, 9110), calnexin (1:1000, Abcam, 22595), atlastin 3 (1:100; Proteintech, 16921-1-AP) and NP (1:1000; gift from Prof Paul Digard); mouse monoclonal against NP (1:1000; Abcam, 20343), virus HA (neat; gift from Prof Paul Digard), M2 (1:500, Abcam, 5416), PDI (1:500, Life Technologies, MA3-019) and Sec31A (1:100; BD Biosciences, 612350); goat polyclonal against ERp57 (1:200; Sicgen, AB0003-200). Secondary antibodies were all from the Alexa Fluor range (1:1000; Life Technologies). Single optical sections were imaged with a Leica SP5 live confocal microscope. Cluster size was quantified as published previously^14^. Distributions (in %) were calculated and plotted by GraphPad Prism. Images were post-processed using Adobe Photoshop CS5 and ImageJ (NIH).

### Live cell imaging

Cells were grown in chambered glass-bottomed dishes (Lab-Tek) and maintained at 37°C in Leibovitz L-15 CO_2_-independent medium (Gibco) during imaging. Samples were imaged using Leica SP5 Inverted or Roper TIRF Spinning Disk (Yokogawa CSU-X1) and post-processed using Adobe Photoshop CS5 and ImageJ (NIH).

For fluorescence recovery after photobleaching (FRAP) analysis, cells were transfected with 250 ng of GFP-NP and immediately superinfected with PR8 at an MOI of 10. At 12 hpi, media was substituted for Leibovitz L-15 media to buffer CO_2_ and data acquisition started on a Roper TIRF Spinning Disk (Yokogawa CSU-X1) with a cage incubator to control temperature at 37 °C. After excitation with a 491 nm laser (Cobolt 491, 100 mW), fluorescence from GFP was detected with a 100x oil immersion objective (Plan Apo 1.49), a bandpass filter (525/45 Chroma), and a photometrics 512 EMCCD camera. All FRAP experiments were performed similarly using iLas FRAP module (Rope Scientific): 2 sec prebleach, 12.18 msec/μm^2^ bleach, 60 sec postbleach at a frame rate of 3 images per second. Bleaching was performed in a variable circular area to target complete viral inclusions. For FRAP analysis, samples were corrected for background fluorescence and acquisition photobleaching as described previously by the Phair method^73^. After normalization, FRAP curves were fitted following the exponential function: Y=Y0 + (Plateau-Y0)*(1-exp(-D*x)), where:

Y0: Y value when X (time) is zero. It is expressed in the same units as Y.

Plateau (must be less than one): Y value at infinite times, expressed in the same units as Y. D:

rate constant, expressed in reciprocal of the X axis time units.

Tau: time constant, expressed in the same units as the X axis. It is computed as the reciprocal of D.

Half-time: time units of the X axis. It is computed as ln(2)/D.

Span (mobile phase): difference between Y0 and Plateau, expressed in the same units as your Y values.

### Tokuyasu – Double Immunogold labeling

Cells infected with PR8, at an MOI of 5, were fixed in suspension using 2% (v/v) formaldehyde (EMS) and 0.2% (v/v) glutaraldehyde (Polysciences) in 0.1 M Phosphate buffer (PB), for 2 h at RT. Subsequently, cells were centrifuged and washed with PB. The aldehydes were quenched using 0.15% (w/v) glycine (VWR) in 0.1 M PB for 10 min at RT. Cells were infiltrated in 12% (w/v) gelatin (Royal) for 30 min at 37°C and centrifuged. The gelatin was solidified on ice, cut into 1 mm^3^ cubes and placed in 2.3 M sucrose (Alfa Aesar) in 0.1 M PB, ON at 4°C. The cubes were mounted onto specimen holders and frozen at -196°C by immersion into liquid nitrogen. Samples were trimmed and cut into 50 nm-thick sections (in a Leica EM-FC7 at -110°C) and laid onto formvar-carbon coated 100-mesh grids.

For immunogold labeling, sections were blocked with PBS/1% BSA for 20 min at RT. Antibody staining was done sequentially in PBS/1% BSA at RT: rabbit anti-GFP (1:500, 1 h), goat anti-rabbit IgG conjugated to 18 nm-gold (1:20, 30 min), mouse anti-NP (1:200, 1 h) and goat anti-mouse IgG conjugated with 6 nm-gold (1:20, 30 min). Gold particles were fixed by applying 1% (v/v) formaldehyde in PBS for 5 min at RT. Blocking and extensive washing were performed in-between stainings. In the final step, gold particles were fixed using 1% (v/v) glutaraldehyde (Polysciences) for 5 min RT. Grids were washed in distilled H_2_O and counterstained using methyl-cellulose–uranyl acetate solution for 5 min on ice. EM images were acquired on a Hitachi H-7650 operating at 100 keV equipped with a XR41M mid mount AMT digital camera. Images were post-processed using Adobe Photoshop CS5 and ImageJ (NIH).

### Correlative light and electron microscopy (CLEM)

Cells, seeded onto gridded dishes (MatTek Corporation, P35G-2-14-C-GRID), were transfected with GFP-NP and simultaneously infected or mock-infected with PR8 at an MOI of 10. At indicated times, cells were fixed, imaged at the confocal microscope Leica SP5 Inverted and finally processed for electron microscopy imaging, as described previously^14^. Sections of 70 nm thickness were cut using a Leica EM-FC7 Ultramicrotome. The regions of interest were acquired with a Hitachi H-7650 operating at 100 keV equipped with a XR41M mid mount AMT digital camera. Images were post-processed using Adobe Photoshop CS5 and ImageJ (NIH).

### Western blotting

Western blotting was performed according to standard procedures and imaged using a LI-COR Biosciences Odyssey near-infrared platform as in^45^. Antibodies used included: rabbit polyclonal against pIRF3 (1:1000; Cell Signal, 4947), virus NP (1:1000), PB1, PB2, PA and NS1 (all at 1:500), kindly provided by Prof. Paul Digard, Roslin Institute, UK; goat polyclonal against green fluorescent protein (GFP) (1:2000; Sicgen, AB0020), GAPDH (1:2000; Sicgen, AB0049) and virus M1 (1:500; Abcam, 20910); mouse polyclonal against virus M2 (1:500; Abcam, 5416). The secondary antibodies used were from IRDye range (1:10000; LI-COR Biosciences).

### Enzyme-linked immunosorbent assay

Detection of IFN-β in the cell supernatants was done using the Verikine^TM^ Human IFN Beta ELISA kit (PBL Assay Science, 41410), range 50-4000 pg/mL, following the manufacturer’s instructions.

### Quantitative real-time reverse-transcription PCR (RT-qPCR)

Extraction of RNA from samples in NZYol (NZYtech, MB18501) was achieved by using the Direct-zol RNA minipreps (Zymo Research, R2052). Reverse transcription (RT) was performed using the transcriptor first strand cDNA kit (Roche, 04896866001). Real-time RT-PCR to detect GAPDH and IFN-β, IFN-α, IL-29 and Viperin was prepared in 384-well, white, thin walled plates (Biorad, HSP3805) by using SYBR Green Supermix (Biorad, 172-5124), 10% (v/v) of cDNA and 0.4 □M of each primer. The reaction was performed on a CFX 384 Touch Real-Time PCR Detection System machine (Biorad), under the following PCR conditions: Cycle 1 (1 repeat): 95°C for 2 min; Cycle 2 (40 repeats): 95°C for 5 s and 60°C for 30 s; Cycle 3: 95°C for 5 s and melt curve 65°C to 95°C (increment 0.05°C each 5 s). Data were analysed using the CFX manager software (Biorad).

Primer sequenced used for real-time RT-qPCR were the following:

GAPDH Fw: 5′-CTCTGCTCCTCCTGTTCGAC-3′;

GAPDH Rv: 5′-ACCAAATCCGTTGACTCCGAC-3′;

IL-29 Fw: 5’-AATTGGGACCTGAGGCTTCT-3’;

IL-29 Rv: 5’- GTGAAGGGGCTGGTCTAGGA-3’;

IFN-β Fw: 5’- CCTGAAGGCCAAGGAGTACA-3’;

IFN-β Rv: 5’- AAGCAATTGTCCAGTCCCAG-3’

IFN-α Fw: 5’- ATGGCCCTGTCCTTTTCTTT-3’

FN-α Rv: 5’- ATTCTTCCCATTTGTGCCAG-3’

Viperin Fw: 5’- TCACTCGCCAGTGCAACTAC-3’

Viperin Rv: 5’- TGGCTCTCCACCTGAAAAGT-3’

## ACKNOWLEDGEMENTS

This project, ALS and FF are supported by Fundação Calouste Gulbenkian-Instituto Gulbenkian de Ciência, Portugal. All other authors are supported by Fundação para a Ciência e a Tecnologia, Portugal: SVC and MA are funded by post-doctoral fellowships: SFRH/BPD/94204/2013 and SFRH/BPD/62982/2009, respectively; TAE is supported by the PhD fellowship PD/BD/128436/2017 and MJA is funded by the FCT investigator contract IF/00899/2013. The authors acknowledge Prof Paul Digard (Roslin Institute, UK) for providing reagents (antibodies, cell lines, viral strains and plasmids), Dr Ron Fouchier (Erasmus, Netherlands) for the reverse genetics plasmids, Dr Colin Adrain (IGC, Portugal) for Sec23 plasmid, and Dr Christoph Dehio for Sec61β-Emerald cell line (University of Basel, Switzerland). The authors thank Dr Fabrice Cordelières from the Bordeaux Imaging Center (INSERM, France), Gabriel Martins (IGC, Portugal), Nuno Pimpão (IGC, Portugal) and Dr Luís Moita (IGC, Portugal) for helpful discussion.

## AUTHOR CONTRIBUTIONS

MJA, MA, SVC designed the experiments; all authors carried out experiments and analyzed the data; MJA supervised the research and conceived the experiments; MJA, MA and SVC wrote the manuscript; all authors contributed to editing the manuscript.

